# Neurocognitive mechanisms of mathematics vocabulary processing in L1 and L2 in South African first graders: An fNIRS study

**DOI:** 10.1101/2025.07.01.662317

**Authors:** Hanrie S. Bezuidenhout, Parvin Nemati, Hadi Borj Khani, Elizabeth Henning, Mojtaba Soltanlou

## Abstract

**Significance:** To learn mathematics, young children require accurate interpretations of mathematics vocabulary. When school language differs from children’s home language, mathematics performance often decreases. Little is known about cortical activation during mathematics vocabulary processing in different languages. This insight will help us to better understand children’s mathematical learning in multilingual societies.

**Aim and approach:** We investigated behavioral and brain responses (fNIRS) of 42 isiZulu and Sesotho (L1) first graders (6.75-7.83 years, 22 girls) who learn mathematics in English (L2) at school when they encounter mathematics vocabulary in L2 compared to L1; and mathematics vocabulary compared to object recognition in L1.

**Results:** The results show that higher accuracy in the L1 mathematics vocabulary, as compared to the L2 mathematics vocabulary, comes with the costs of higher cognitive demands in the right superior and middle frontal gyri for first graders. Mathematics vocabulary required longer response time than object recognition and a higher activation in the right superior frontal gyrus. No parietal difference was observed between conditions.

**Conclusions:** First graders with no automatization of mathematics vocabulary processing, still demand frontal cognitive resources. This study is a good example of how educational neuroimaging compliments our interpretation of behavioral outcomes and environmental factors such as multilingualism.

## 1 Introduction

Mathematics learning requires, among other skills, an accurate interpretation of mathematics vocabulary. It is children’s understanding of key words, phrases and abbreviations that are often used in mathematics instruction, assessments and discussions^1,2,3,4^. Interpreting mathematics vocabulary is more challenging for children who learn mathematics in a different language (L2) than their home language (Ll). Unfamiliar technical terms (e.g. decrease), terms with more than one meaning (e.g. difference) and homonyms (e.g. sum) are examples that can make interpretation, understanding and reasoning more difficult in L2^5^. This is also the case for children whose home language differs from their school language, which is common in multilingual societies like South Africa. For instance, Bezuidenhout^6^ (2019) found that South African first graders who receive mathematics instruction in L2, achieved higher scores on mathematics and mathematics vocabulary assessments in L1 (isiZulu/Sesotho) than in L2 (English), with concurrent and predictive associations between mathematics vocabulary and mathematics performance. She concluded that the children in that sample had more exposure to L1 mathematics vocabulary in their first six years and that the limited understanding of L2, their school language, directly influenced their lower performance in L2. Because mathematics vocabulary predicts mathematics performance throughout children’s school achievement^7,8^ the development of mathematics vocabulary is crucial in the early development years. Therefore, it is crucial to understand the development of mathematics vocabulary and its underlying neural mechanisms in young children, particularly in multilingual societies.

The understanding and interpretation of mathematics vocabulary in the context of general language structures is inherently needed for mathematical thinking^1,9^. For instance, mathematics tasks are often couched in word problems and require an understanding of both general language and vocabulary specific to mathematics^5^. Vygotsky^10^ (1968) was one of the first theorists to argue that language is required to reason logically and to represent and process concepts^11^. The idea that language supports mathematics concept development in this way^12,13^, is validated by well established links between mathematics and language abilities^9,14,15,16^. To explain these links, Gentner and Goldin-Meadow^17^ (2003) propose that language is like a lens that enables us to look at concepts; and is also like a tool that enlarges one’s existing representations of the world. Gopnik and Melzoff^18^ (1997) proposed a third metaphor by arguing that language is also an input for learning concepts (see also Levine & Baillargeon^13^, 2016). This means that young children who are more often exposed to everyday mathematical discussions with caregivers, parents and teachers, are more likely to understand mathematics concepts than those with limited mathematical linguistic input. In this study, we focus on one specific component of language that has repeatedly been shown to contribute to the development of mathematics concepts^1,19,20,21^, namely mathematics vocabulary. Children without mathematics vocabulary comprehension are unable to follow mathematics instruction or explain their reasoning.

Children with an elaborate mathematical vocabulary are better equipped to engage in advanced mathematics activities, such as comparison of numbers, descriptions of relationships or properties of numbers or shapes or describing mathematical procedures. For this reason, several researchers^19,22,23^ have highlighted the importance of frequent exposure to the correct use of mathematics vocabulary in the early years. This poses a challenge to children who are less frequently exposed to L2 mathematics vocabulary during preschool years and transition to L2 as a language of learning in the first grade. Because young L2 learners may have a limited “linguistic toolbox” of mathematics vocabulary or look through a “blurry linguistic lens”, they may struggle to understand mathematics concepts. Thus, even though young L2 children may conceptually comprehend mathematics concepts and be able to demonstrate it in L1, their limited understanding of L2 may limit their mathematics performance when assessed in L2. While behavioral research^21,24^ focus on developing intervention programs for improving mathematics vocabulary, a better understanding of the underlying cognitive mechanisms of mathematics vocabulary processing, especially in L2, allows both teachers and research practitioners to tailor pedagogical tools.

Limited neuroimaging studies have compared differences in brain activation between L1 and L2 mathematics processing. Wang et al.^25^ (2007) compared brain activation differences in L1 and L2 processing of mathematics tasks (i.e. numerical concept processing and exact calculation of multiplication and addition). They conducted an fMRI study on young adults to describe the difference in underlying neural mechanisms of mathematics processing in Mandarin (L1) compared to English (L2). The results showed that calculation tasks performed in L2 involved higher neural activation in the left hemisphere, including the inferior frontal gyrus (Broca’s area, which is responsible for language processing), compared to L1. The participants in their study translated or transcoded L2 questions into L1 to perform computation tasks in more familiar vocabulary and language structures. Their findings suggest that young adults process L2 calculation tasks through the L1 language system because the arithmetic memory system is not well developed in bilinguals’ L2. Typically, mathematics is processed in the frontoparietal network^26,27^. Different parts of the parietal regions including the bilateral intraparietal sulcus (IPS) for quantity representation, the left angular gyrus (AG) for language-mediated processes during mathematics tasks (e.g. verbal short-term memory^27^) and superior parietal lobule for spatially-mediated processes are involved in mathematical processing. Additionally, the prefrontal cortex, related to executive functions, serves a supportive domain-general role^27^. In a functional near infrared spectroscopy (fNIRS) study, Sugiura et al.^28^ (2011) found higher cortical activation in the bilateral superior/middle temporal and inferior parietal areas in 6- to 10-year olds when processing general auditory L1 vocabulary than L2 vocabulary. They suggest that greater neural activation in L1 than L2 means that young children more easily process the phonology of L1, while phonologically unfamiliar L2 words are likely processed like nonword auditory stimuli. An outstanding question is how young children, as they start compulsory school, process mathematics vocabulary in L1 and L2. Is higher cortical activation expected in L2 like the adults in Wang et al.’s study or higher activation in L1 like the children in Sugiura’s study?

Draper et al.^29^ (2022) highlighted the underrepresentation of studies in majority countries such as sub-Saharan Africa and calls for a better understanding of cognitive development in countries with diverse cultural and social characteristics, including multilingualism. South Africa is a multilingual country with 12 official languages. Most children receive mathematics instruction in L1 until the third grade and then transition to English as language of instruction in the fourth grade. However, several schools have a custom designed language policy and children already receive tuition in English from the first grade, with some code switching to L1 and translanguaging practices. During code-switching^30,31^, children preserve linguistic features of languages^32^ while switching between two defined linguistic structures within a sentence or by repeating sentences or words in a second language. Translanguaging^33,34,35^, on the other hand, dismantles distinct language features of named languages by spontaneously integrating spoken language into a single system. For instance, isiZulu and Sesotho speakers in South Africa code-switch and practice translanguaging by using English number words in their isiZulu or Sesotho sentences. The other outstanding question is how young children in a multilingual society, like South Africa, process mathematics vocabulary in L1 and L2.

## 2 The current study

To address the above outstanding questions, the current study attempts to provide insight about the underlying neural mechanisms of mathematics vocabulary in multilingual children in the majority world^36^. In the present within-subject study, we investigated the neural network activation differences of L1 (Sesotho/isiZulu – two of the official and most frequently used languages of South Africa) and L2 (English) during mathematics vocabulary tasks in first graders in South Africa. We aimed to understand i) the brain responses in L1 and L2 processing of mathematics vocabulary and ii) brain activation during L1 processing of mathematics vocabulary and non-mathematics vocabulary or object recognition. For mathematics vocabulary, children compared two sets of objects to determine where it is more or less and used keypress to indicate their answer. Prerecorded auditory instructions were given, e.g. “where is more/less”. For object recognition, children had to recognize specific animals, e.g. “where is the bird?” and also used keypress to answer. We investigated brain responses in the bilateral frontoparietal network using functional near-infrared spectroscopy (fNIRS). It is a non-invasive brain imaging technique that allows studying brain activity in naturalistic settings^37,38^. Additionally, fNIRS is safe, cost-effective, and suitable for use with diverse populations, including young children^39^.

The research question that directed the study was: What are the differences and similarities in brain responses of first graders in multilingual settings when they encounter i) mathematics vocabulary in L2 compared to L1; and ii) mathematics vocabulary compared to object recognition in L1? Our first hypothesis built on the findings of Bezuidenhout^40^ (2021) that South African first graders achieve lower mathematics vocabulary scores in L2 than in L1^7^. Based on their lower performance, we also expected higher cognitive engagement of the supporting prefrontal brain areas^25^ for L2 processing, as compared to L1. Secondly, we expected slower and less accurate responses during mathematics vocabulary than object recognition in L1, along with higher brain activation in the frontoparietal network.

## 3 Materials and methods

### 3.1 Participants

In a similar, though behavioral study, Bezuidenhout^7^ (2018) reported a large Cohen’s *d* effect size of 0.92. Based on Bezuidenhout^7^ (2018), we calculated the sample size by using G*Power for paired *t*-tests (two tailed, alpha = 0.05, power = 0.9), which suggested a total of 12 children. However, to account for smaller effect sizes in fNIRS studies, we used a medium Cohen’s *d* effect size of 0.5, which suggested a total of 34 children for the current within-subject design.

Altogether, 42 children participated in the current study. Children would have been excluded from the study if they had learning difficulties, a diagnosed neurological or mental illness, diagnosed diabetes or blood pressure (hypertension), uncorrected diagnosed visual or auditory impairment or had a serious head injury, such as concussion, or if they did not finish the study. There were no such children. One of the challenges of an fNIRS study in sub-Saharan Africa is that some participants may have thick, dark hair or wear braids. Therefore, we asked them not to braid their hair when participating in the study.

The study was approved by the Research Committee of the University of Johannesburg which oversees research projects being conducted at the school, as well as the Ethics Committee of the Education Faculty of the University of Johannesburg (Sem 1-2022-035). The study was conducted according to the ethical guidelines and principles of the international Declaration of Helsinki. Children received incentives such as toys and stickers for their participation.

The study was conducted at the Cognition Lab at the University of Johannesburg which is located next to the children’ school. The school was purposefully chosen to investigate the effect of learning mathematics in L2 in young children who have not been exposed to English as frequently as older children. When children’s school language differs from their home language, their mathematics performance often decreases. The isiZulu- and Sesotho-speaking first graders learn mathematics in English (L2) at this school. With enrolment to this teaching school, caregivers consent to their children’s participation in research experiments. However, this is one of the first neuroimaging studies that were conducted and therefore we also obtained written informed consent from the caregivers and participating children. The school is a quintile 2 school which means that there are no school fees payable. These quintile poverty rankings, with quintile 1 being the lowest, are determined nationally according to the income, literacy and unemployment levels of the community around the school, as well as certain infrastructural factors such as permanent vs. temporary classes, classroom equipment, access to water and sanitation and safety measures^41^.

### 3.2 Procedure

In this within-subject study, we measured brain activation as well as six behavioral tasks to assess children’s general cognitive skills, of which two assessments were completed in both L1 and L2 for comparison purposes. Children whose caregivers consented to participation were collected from the school (which is on the premises of the university close to the lab) in groups of three to four children at a time and returned to their classes after participation. The experimenter introduced the children to the lab and explained the study. While one child engaged in the behavioral assessments in a separate, another completed the neuroimaging tasks. The remaining children were given toys in an adjacent room with a caregiver while waiting for their turn to participate.

### 3.3 Behavioral assessments

#### 3.3.1 Mathematics language

The Mathematics Vocabulary Test (MVT)^42^ is a 24-item test for preschool and early grades children. It assesses numerical language qualifiers (more, many, just as many, fewer, few), comparative language (same size, bigger, tallest, biggest, big/large, tall, shortest, small, smaller, short) and spatial language (in between, first, last, on top of, behind, above, under, after, in front of). Each item requires the child to point to the picture that describes a specific concept. For instance: “Put your finger on the picture with more bugs”. None of the items requires exact quantitative knowledge, but only approximate numerical knowledge. One point is awarded for each correct response and scores are calculated based on the total number of correct responses out of 24. Mathematics vocabulary was assessed in L1 (Sesotho/ isiZulu) and L2 (English) to compare the children’s performance in their home language and language of instruction.

#### 3.3.2 Mathematics

The Preschool Early Numeracy Skills Test (PENS)^43^ is a 25-item numeracy test that assesses numeracy skills of children between three and six years. The items assess set comparison, numeral comparison, one-to-one correspondence, number order, identifying numerals, ordinality, and number combinations. Some items are multiple choice and others are free response. Except for the first question where a score out of five is determined, one point is awarded for each correct response and scores are calculated based on the total number of correct responses out of 29. The test stops if the child answers three consecutive questions incorrectly. Mathematics performance was assessed in L1.

#### 3.3.3 Verbal short-term memory

The digit span task as a subtest of the Wechsler Intelligence Scale for Children (WISC-IV) was used to assess how well children can recall spoken sequences of consonant digits (one digit per second). The test starts with sequences of two digits, which are increased by one if the child correctly recalls at least two out of three sequences. Each correct answer scored one point and the total score is the sum of all points. Only forward recalls were included because of the children’s young age. Digit span was conducted in L1 and L2 to compare responses between the two languages.

#### 3.3.4 Nonverbal IQ

Subtest 3 of the Culture Fair Test (CFT)^44^ is a 15-item pencil and paper subtest that assesses similarity recognition of 4- to 8-year-old children. Children identify as many similarities as they can, with a maximum of 15, within 90 seconds. For each item, one picture is presented with five similar pictures. The child sees a picture on the left side of a table and then selects one out of five pictures –the one which is exactly the same as the first picture. One point is awarded for each correct response and scores are calculated based on the total number of correct responses out of 15. Instructions were given in L1.

#### 3.3.5 Listening comprehension

Gogo’s Dog (grandmother’s dog)^45^ with a listening comprehension scale^46^ based on the Shell–K listening comprehension protocol^47^ assesses listening comprehension. The test consists of one story (fiction), with 15 questions, ranging in difficulty from basic factual questions to questions, which require the child to infer answers from the text. The child responds orally while the tester writes down their answers. This is not a timed test. One point is awarded for each correct response and scores are calculated based on the total number of correct responses out of 15. Listening comprehension was assessed in L1.

#### 3.3.6 Mathematics Anxiety

A modified version of the Abbreviated Mathematics Anxiety Scale (AMAS)^48^ is a 9-item scale that assesses mathematics anxiety. For each item the child needs to indicate whether he or she experiences low (1), some (2), moderate (3), quite a bit (4) or high (5) anxiety. Because first graders are not yet able to assign numbers one to five to their level of anxiety in a given situation, the child was presented with five different faces, representing numbers one to five. A total score is calculated by adding scores of 9 items. Mathematics anxiety was evaluated in L1.

### 3.4 fNIRS tasks

During the neuroimaging tasks, the child wore a fNIRS cap which was carefully fitted onto the head. While calibration took place, the fNIRS cap was covered with an additional black cap to minimize light exposure. The experiment was conducted in a light attenuated room, where all the lights were switched off before starting the calibration and experiment.

The neuroimaging experiment consisted of two tasks: an experimental task containing 12 blocks of mathematics vocabulary processing, including six blocks in L1 and six blocks in L2; and a control task in L1, containing six blocks of object recognition (see Fig. 1). To balance our design, we also had a control condition in L2, but we did not include it in the analysis as it was not relevant to our research questions. Within the experimental tasks, the order of the L1 and L2 blocks were counterbalanced across children. Each experimental and control task started with two practice blocks and each block consisted of 6 trials. After completing the practice blocks, the experimenter verbally checked with the children to make sure that they understood the instructions. The experimenters were three trained native isiZulu and Sesotho speakers, and verbal instructions were given in the children’s L1.

**Fig. 1.**
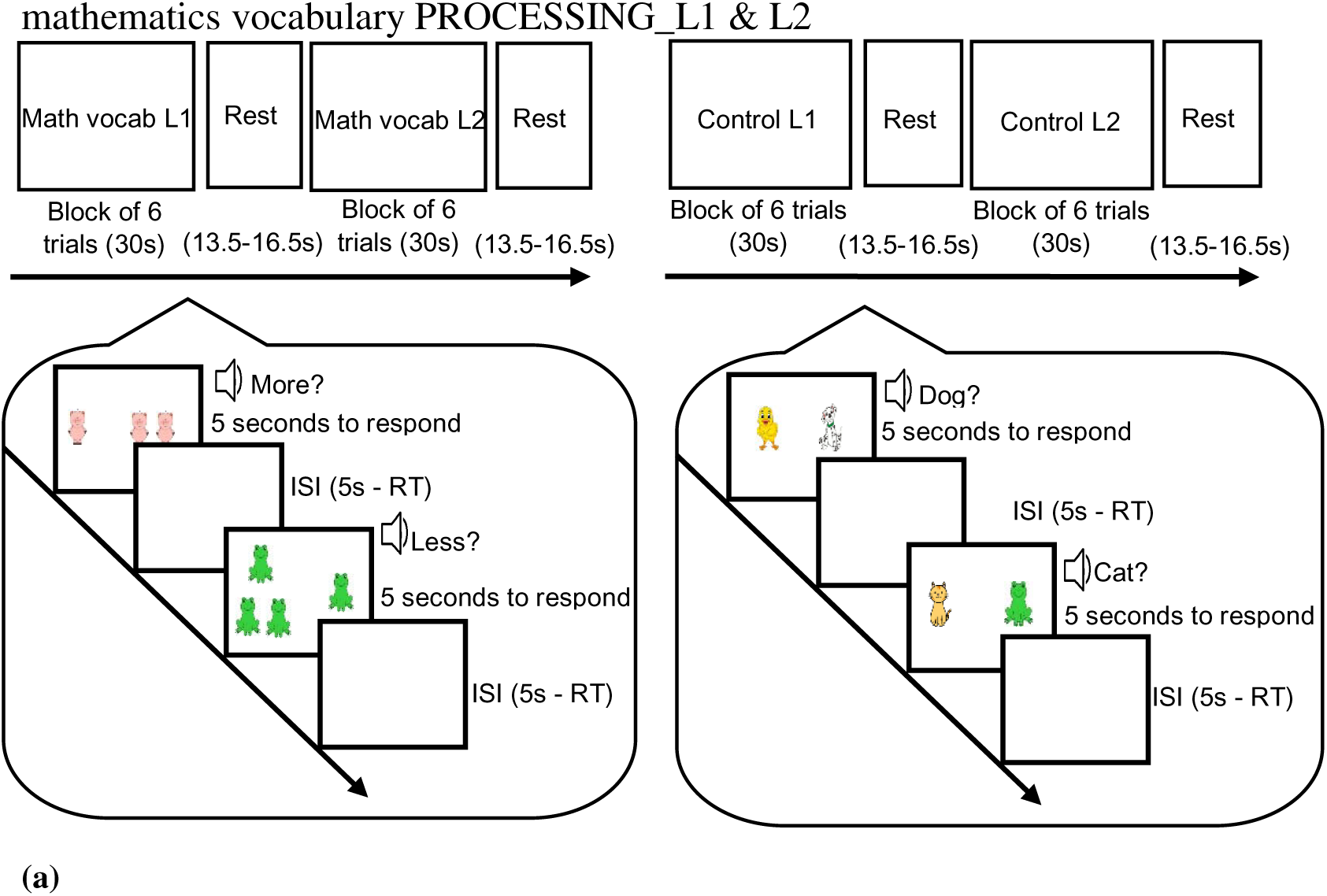
(a) Experimental tasks (i.e. mathematics vocabulary) and control tasks (i.e. object recognition) were presented in a mixed block design in L1 and L2. Participants selected the correct picture corresponding to the auditory input with a key press. Note: ISI: inter-stimulus interval; RT: Response Time.

For each experimental trial, children saw two sets of pictures side-by-side on a computer screen, while hearing one of two words randomly: *more* or *less*. In half of the trials, they had to identify more items, while in the other half, they had to identify less items. By using keypress, children then chose the set of pictures related to the word they heard (see Fig. 1). They pressed key A for selecting the left set and key L for selecting the right set. For example, if one pig appeared on the left side of the screen and two pigs on the right, while the child heard “more”, the child had to press key L. Sets included one, two, three or four animals, which was within the subitizing range^49^. For each control trial, children saw two different, but the same number of animals on the screen, heard the name of one of the animals on the screen and selected the matching picture by using keypress. In half of the trials, they had to select the left one, while in the other half, they had to select the right one. They needed to press key A for selecting the left one and key L for selecting the right one. Each trial was presented for a maximum of five seconds in both experimental and control tasks. If the child responded faster than 5 seconds, a blank screen appeared for the remainder of the 5 seconds before the next trial appeared, as the inter-stimulus interval (ISI). Between each block, there was a resting time of 13.5-16.5 seconds with a jittering step of 0.5 seconds, which was randomly selected from a pool of seven possible lengths. Although feedback was given during the practice blocks, no feedback was given during the experimental questions. The neuroimaging experiment was conducted by using the program OpenSesame^50^.

## 4 fNIRS Recording, Preprocessing, and Analysis

A multichannel fNIRS device, the NIRScout system (NIRx Medical Technologies, LLC), was used to monitor and record cortical hemodynamic responses during the mathematics vocabulary and the control tasks. There were 15 emitters that emit light at wavelengths between 760 and 850 nm, and 11 Avalanche Photodiode detectors. A sampling frequency of 4.1667 Hz and an average distance of 3 cm between optodes were used for data acquisition.

At the beginning of the data processing, a rigorous quality assessment and control procedure was implemented. The QT-NIRS toolbox^51,52^, which is available on the QT-NIRS GitHub (https://github.com/lpollonini/qt-nirs), was used to identify and exclude channels with inadequate data quality. Following Fonseca et al.^36^ (preprint), channels below thresholds (SCI threshold = 0.5, Q threshold = 0.3, PSP threshold = 0.1) were excluded from subsequent analyses (Fig.2a).

**Fig. 2.**
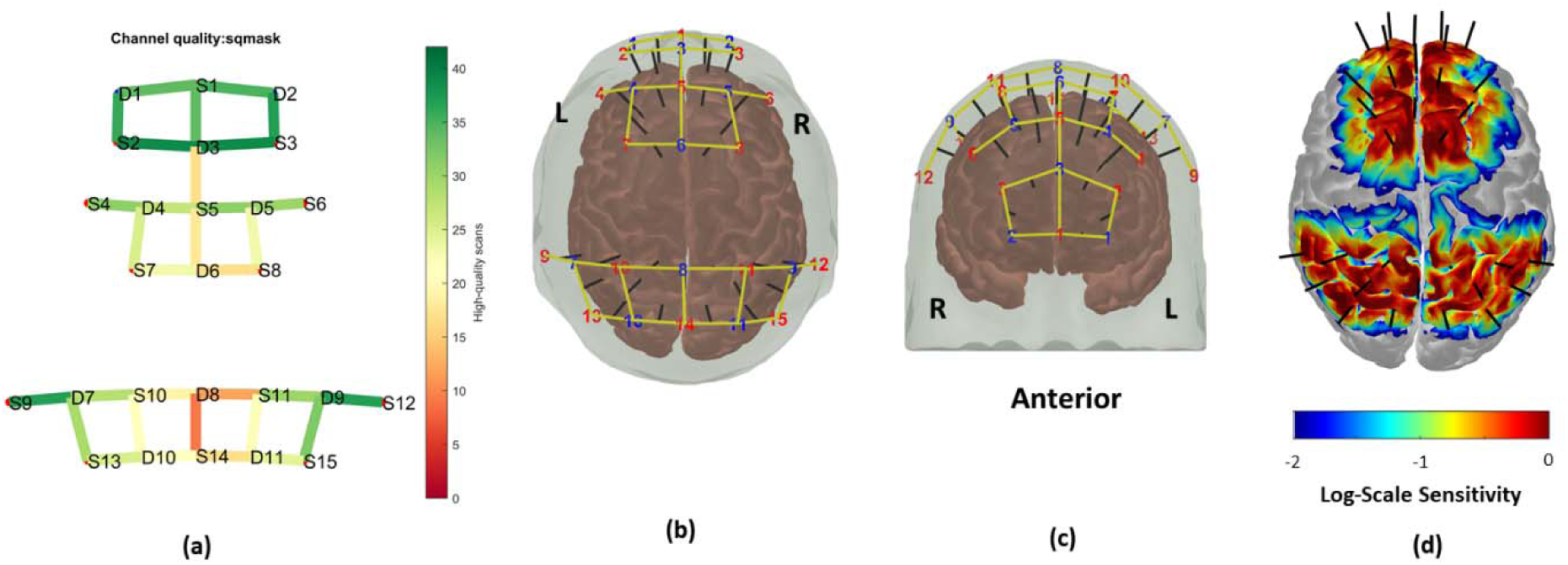
(a) The colour bar indicates a signal quality map illustrating the number of participants per channel with a quality index exceeding 30%. (b) Superior view and (c) anterior view of the head model showing the positions of each source and detector, along with the projection of channels onto the cortical surface, highlighting the corresponding measurement locations and cortical regions. They show the positions of sources (red) and detectors (blue) distributed across the scalp. Yellow lines represent the channels formed by source-detector pairs, with black markers indicating the projections of these channels onto the cortical surface. (d) Sensitivity analysis based on the forward model simulation performed using AtlasViewer, illustrating the spatial sensitivity of the optodes to underlying cortical regions and the interaction of light propagation with brain tissue.

In Hernandez & Pollonini^51^ (2020), these quality control steps were followed to retain channels that had acceptable quality metrics (QI > 30%, Cohen’s d > 0.5). We have estimated the MNI coordinates for the retained channel numbers and source-detector pairs using AtlasViewer^53^ in Table 1. Fig. 2(b&c) illustrates the positions of each source and detector on the head model, presented from both superior and anterior views. The figure also includes the projection of each channel onto the cortical surface, providing a clear visualization of the measurement locations and their corresponding cortical regions. Fig. 2(d) also shows the sensitivity analysis derived from the forward model simulation using AtlasViewer. This analysis quantifies the spatial sensitivity of the optodes to underlying cortical regions, providing a detailed visualization of how the light propagation interacts with the brain tissue and contributes to the measurement signals.

**Table 1.** Channel Registration using AtlasViewer.

| Ch | Src | Det | Ch Coord (Monte Carlo) | Ch Coord (MNI) | Area |
| --- | --- | --- | --- | --- | --- |
| 1 | 1 | 1 | 144 149 214 | -12 72 1 | L medial orbital frontal |
| 2 | 1 | 2 | 117 150 212 | 15 70 0 | R medial orbital frontal |
| 3 | 1 | 3 | 132 136 197 | 0 55 14 | superior medial frontal |
| 4 | 2 | 1 | 151 144 202 | -19 60 6 | L superior frontal |
| 5 | 2 | 3 | 143 128 200 | -11 58 22 | L superior frontal |
| 6 | 3 | 2 | 108 140 200 | 24 58 10 | R superior frontal |
| 7 | 3 | 3 | 117 126 202 | 15 60 24 | R superior frontal |
| 8 | 5 | 3 | 130 108 199 | 2 57 42 | superior medial frontal |
| 9 | 4 | 4 | 151 114 183 | -19 41 36 | L middle frontal |
| 10 | 5 | 4 | 141 96 191 | -9 49 54 | L superior frontal |
| 11 | 7 | 4 | 152 91 175 | -20 33 59 | L superior frontal |
| 12 | 5 | 5 | 119 96 190 | 13 48 54 | R superior frontal |
| 13 | 6 | 5 | 107 113 180 | 25 38 37 | R middle frontal |
| 14 | 8 | 5 | 117 104 164 | 15 22 46 | R superior frontal |
| 15 | 5 | 6 | 131 84 175 | 1 33 66 | supplementary motor area |
| 16 | 7 | 6 | 141 90 158 | -9 16 60 | L supplementary motor area |
| 17 | 8 | 6 | 118 77 162 | 14 20 73 | R supplementary motor area |
| 18 | 9 | 7 | 191 103 102 | -59 -40 47 | L inferior parietal |
| 19 | 10 | 7 | 164 92 103 | -32 -39 58 | L superior parietal |
| 20 | 10 | 8 | 144 72 99 | -12 -43 78 | L postcentral |
| 21 | 11 | 8 | 113 71 95 | 19 -47 79 | R postcentral |
| 22 | 11 | 9 | 88 82 97 | 44 -45 68 | R superior parietal |
| 23 | 12 | 9 | 73 105 98 | 59 -44 45 | R inferior parietal |
| 24 | 10 | 10 | 146 92 97 | -14 -45 58 | L superior parietal |
| 25 | 11 | 11 | 99 78 86 | 33 -56 72 | R superior parietal |
| 26 | 13 | 7 | 160 111 96 | -28 -46 39 | L inferior parietal |
| 27 | 13 | 10 | 155 110 86 | -23 -56 40 | L inferior parietal |
| 28 | 14 | 8 | 129 87 97 | 3 -45 63 | precuneus |
| 29 | 14 | 10 | 142 84 75 | -10 -67 66 | L precuneus |
| 30 | 14 | 11 | 116 85 72 | 16 -70 65 | R precuneus |
| 31 | 15 | 9 | 88 102 89 | 44 -53 48 | R inferior parietal |
| 32 | 15 | 11 | 93 95 72 | 39 -70 55 | R inferior parietal |

Note: R: right, L: left

To mitigate direct current shifts, baseline corrections were applied to the raw intensity signals (SD = 3). After applying the Temporal Derivative Distribution Repair (TDDR) method to address motion artifacts and enhance data robustness^54^, a bandpass filter of 0.01–0.2 Hz was applied^55^. This filtering step was implemented to isolate and retain the frequency components corresponding to typical hemodynamic responses while removing low-frequency drifts and high-frequency noise. To further improve signal fidelity, minimize artifacts, and identify regressors, the AR-IRLS regression model^56^ was employed using the BrainAnalyzIR toolbox^56^, as one of the most commonly used analysis tool^57^.

For the statistical analysis, a group-level approach was employed to investigate the effects of the conditions: mathematics vocabulary in L1 (L1Num), mathematics vocabulary in L2 (L2Num), and object recognition control condition in L1 (L1Con), as well as their contrasts, including L1Num–L2Num and L1Num–L1Con for each channel. A general linear mixed regression model was utilized to compute the beta coefficients for oxyhaemoglobin (HbO) and deoxyhaemoglobin (HbR) corresponding to each condition and contrast. To address the issue of multiple comparisons, the Benjamini-Hochberg False Discovery Rate (FDR) correction (pFDR) was applied to reduce false positivity. The analysis focused on identifying group-level effects across the conditions and their contrasts to provide insights into the distinct patterns of brain activation associated with each condition and contrast. The different brain regions were identified as activated based on their respective channels (source-detector pairs) with significant t-values (activation strength) and q-values (statistical significance). A significant increase/higher HbO and or a significant decrease/lower HbR are interpreted as increase/higher brain activation in the corresponding channels.

## 5 Behavioral Analyses

Eleven and 12 children were excluded from the accuracy and RT analyses of mathematics vocabulary presented in L1 and L2, respectively, due to technical problems with the fNIRS recording (see Table 2). There were also some missing data for some demographic information (i.e, age, sex, and siblings, see Table 2). Mean scores for all measures are reported in Table 2 and the differences between L1 and L2 for the mathematics language and digit span tasks were tested using paired *t*-tests.

**Table 2.** Descriptive Statistics for Study Variables.

| Measures | *n* | *M* | *SD* | Min - Max |
| --- | --- | --- | --- | --- |
| Age | 37 | 7.19 | 0.30 | 6.75 - 7.83 |
| Sex (Female) | 22 |  |  |  |
| Siblings | 37 | 1.62 | 0.80 | 1 - 4 |
| Nonverbal IQ | 42 | 8.95 | 2.58 | 2 - 14 |
| Listening Comprehension | 42 | 7.83 | 2.66 | 3 - 13 |
| Math Anxiety | 42 | 25.80 | 4.00 | 18 - 38 |
| Mathematics | 42 | 21.00 | 5.62 | 9 - 28 |
| **Language 1** |  |  |  |  |
| Mathematics Language | 42 | 21.10 | 3.05 | 18 - 30 |
| Digit Span Forward | 42 | 3.00 | 0.80 | 2 - 5 |
| Accuracy in Experimental Condition | 31 | 0.86 | 0.20 | 0.36 - 1.00 |
| Accuracy in Control Condition | 31 | 0.86 | 0.13 | 0.50 - 0.95 |
| RT in Experimental Condition | 31 | 1836.20 | 355 | 1399 - 2620 |
| RT in Control Condition | 31 | 1736 | 348 | 1208 - 2684 |
| **Language 2** |  |  |  |  |
| Mathematics Language | 42 | 21.20 | 3.54 | 15 - 29 |
| Digit Span Forward | 42 | 3.55 | 1.06 | 2 - 6 |
| Accuracy in Experimental Condition | 30 | 0.82 | 0.20 | 0.43 - 1.00 |
| RT in Experimental Condition | 30 | 1780 | 387 | 1175 - 2480 |

Note. Bold *p*-values depict $p < .05$
*6.2 Mathematics vocabulary in L1*

To test the behavioral performance for the first hypothesis (i.e., children respond slower and less accurately when mathematics vocabulary is presented in L2 than in L1), we conducted paired *t*-tests to compare response times (RT) and accuracy. Cohen’s *d* was used to report effect sizes and can be interpreted as *d* = 0.2 indicating small, *d* = 0.5 medium, and *d* = 0.8 large effect^58^. To test the behavioral performance for the second hypothesis (i.e., children respond slower and less accurately to mathematics vocabulary in L1 than the control in L1), we conducted paired *t*-tests to compare RT and accuracy.

Correlation Analysis was used to test the relationships between study measures (i.e., mathematics vocabulary, mathematics language, mathematics, verbal short-term memory, nonverbal IQ, listening comprehension and mathematics anxiety) together with demographic factors. All the analyses were conducted by Jamovi (Version 2.5, 2024).

## 6 Results

### 6.1 Descriptive Statistics

Children demonstrated variability in performance across conditions, as reported in Table 2.

### 6.2 Mathematics vocabulary in L1

The brain activation significantly increased in the right superior frontal gyrus, bilateral middle frontal gyri, bilateral inferior parietal gyri, the left superior parietal gyrus, and the right precuneus during mathematics vocabulary in L1 (Table 3 and Fig. 3).

**Fig. 3.**
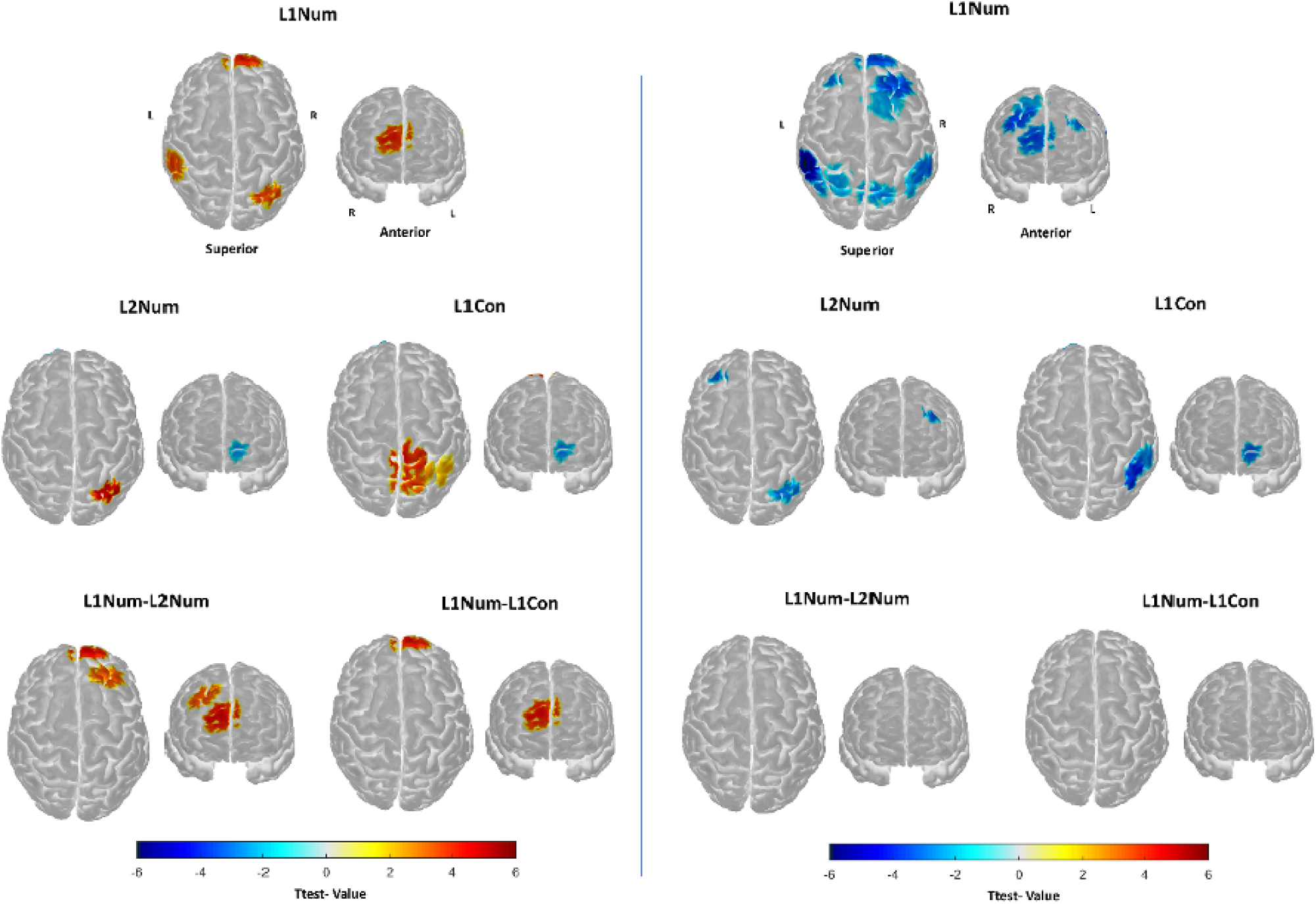
Spatial distribution of task-related brain activation for mathematical vocabulary in L1 (L1Num), mathematical vocabulary in L2 (L2Num), non-mathematical control in L1 (L1Con), and their contrasts (L1Num-L2Num, L1Num-L1Con). The left panel depicts regions with changes in HbO concentration, while the right panel shows regions with changes in HbR concentration. Each brain map highlights significant cortical regions associated with the respective tasks and contrasts, overlaid on a standard brain template. Warm colors (yellow to red) represent positive t-values, while cool colors (blues) indicate negative t-values. The scale bar denotes t-values ranging from -6 to 6.

**Table 3.**
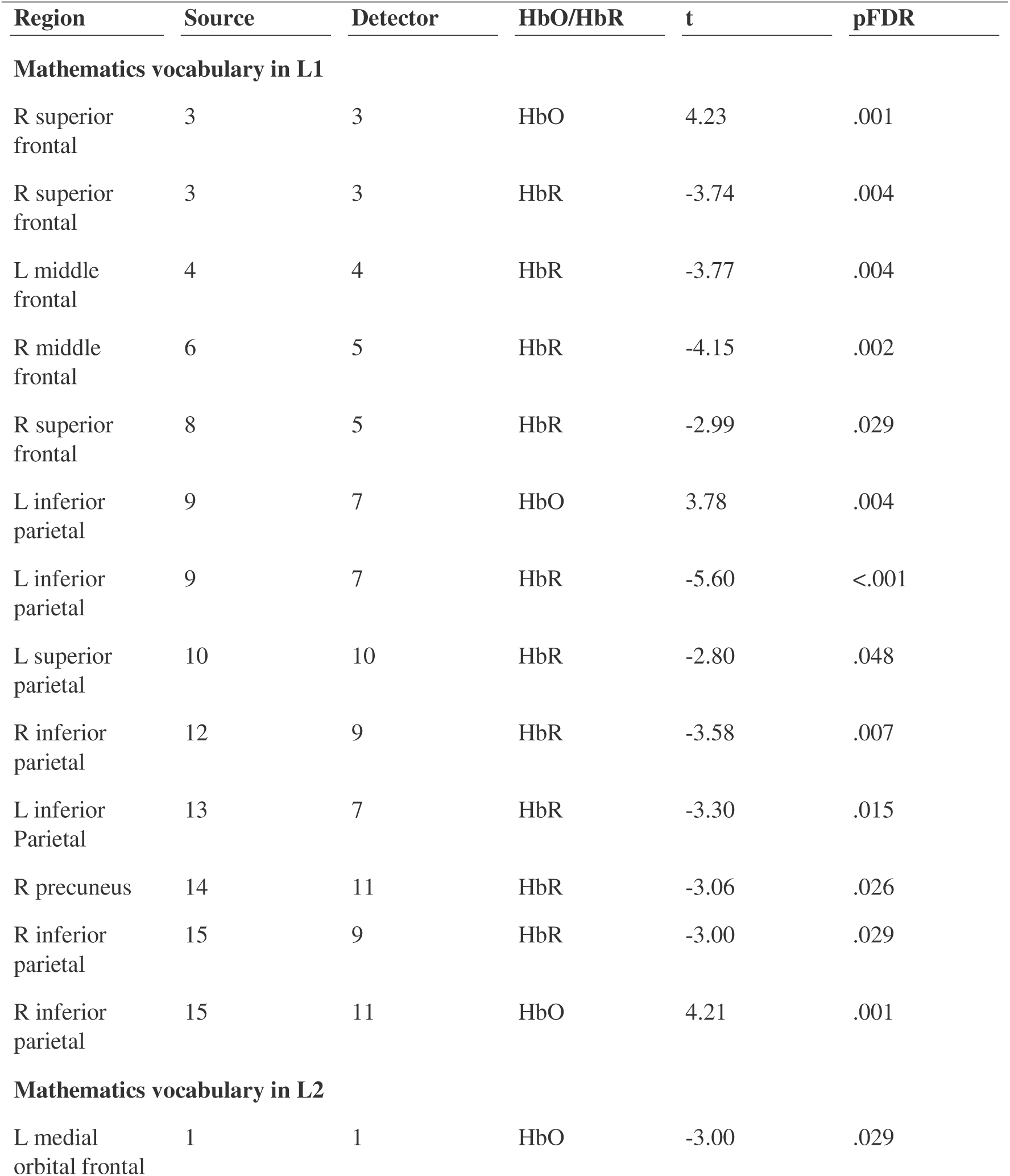
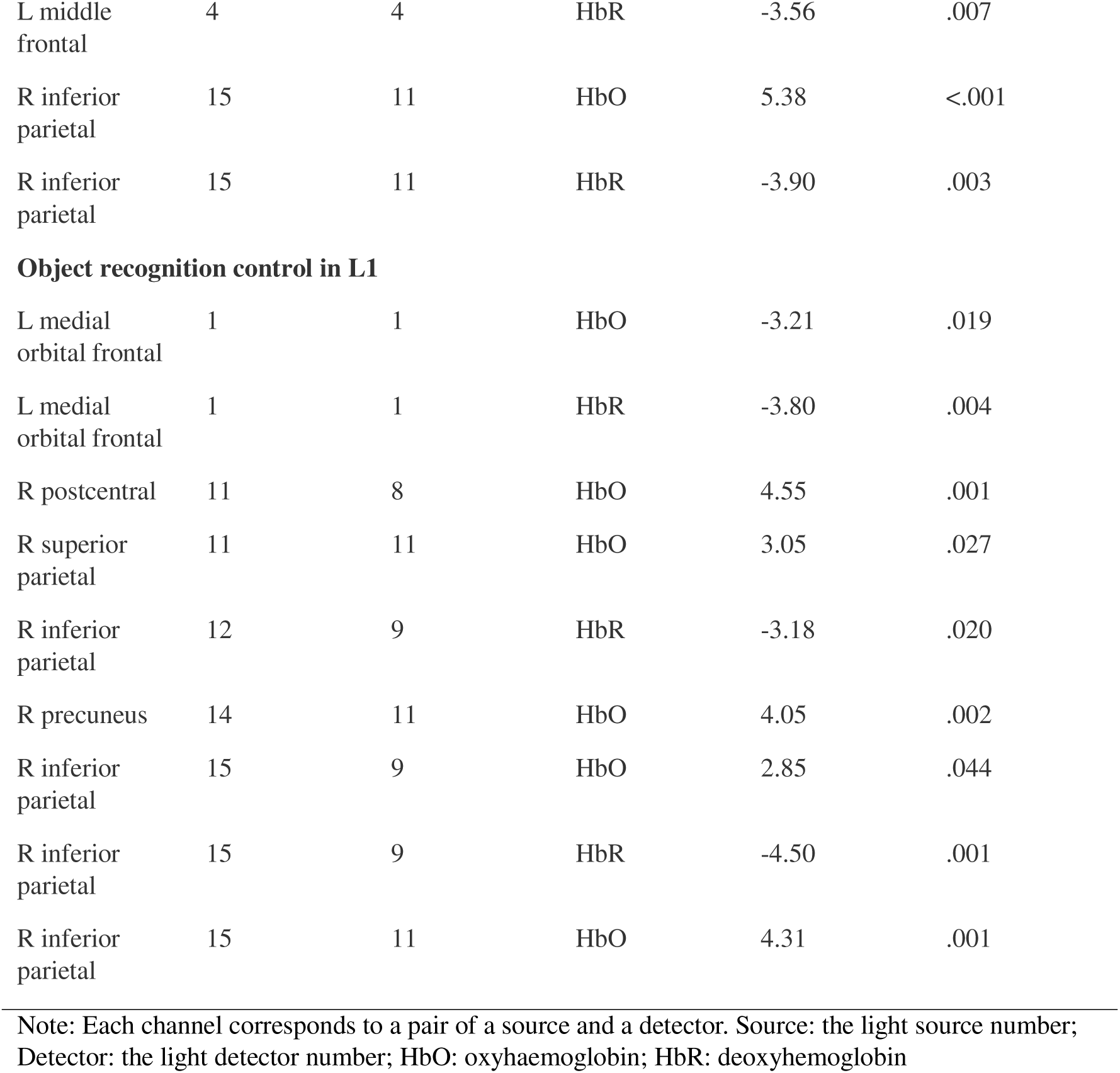
Significant channels in the three different conditions.

| Region | Source | Detector | HbO/HbR | t | pFDR |
| --- | --- | --- | --- | --- | --- |
| **Mathematics vocabulary in L1** |  |  |  |  |  |
| R superior frontal | 3 | 3 | HbO | 4.23 | .001 |
| R superior frontal | 3 | 3 | HbR | -3.74 | .004 |
| L middle frontal | 4 | 4 | HbR | -3.77 | .004 |
| R middle frontal | 6 | 5 | HbR | -4.15 | .002 |
| R superior frontal | 8 | 5 | HbR | -2.99 | .029 |
| L inferior parietal | 9 | 7 | HbO | 3.78 | .004 |
| L inferior parietal | 9 | 7 | HbR | -5.60 | <.001 |
| L superior parietal | 10 | 10 | HbR | -2.80 | .048 |
| R inferior parietal | 12 | 9 | HbR | -3.58 | .007 |
| L inferior Parietal | 13 | 7 | HbR | -3.30 | .015 |
| R precuneus | 14 | 11 | HbR | -3.06 | .026 |
| R inferior parietal | 15 | 9 | HbR | -3.00 | .029 |
| R inferior parietal | 15 | 11 | HbO | 4.21 | .001 |
| **Mathematics vocabulary in L2** |  |  |  |  |  |
| L medial orbital frontal | 1 | 1 | HbO | -3.00 | .029 |
| L middle frontal | 4 | 4 | HbR | -3.56 | .007 |
| R inferior parietal | 15 | 11 | HbO | 5.38 | <.001 |
| R inferior parietal | 15 | 11 | HbR | -3.90 | .003 |
| **Object recognition control in L1** |  |  |  |  |  |
| L medial orbital frontal | 1 | 1 | HbO | -3.21 | .019 |
| L medial orbital frontal | 1 | 1 | HbR | -3.80 | .004 |
| R postcentral | 11 | 8 | HbO | 4.55 | .001 |
| R superior parietal | 11 | 11 | HbO | 3.05 | .027 |
| R inferior parietal | 12 | 9 | HbR | -3.18 | .020 |
| R precuneus | 14 | 11 | HbO | 4.05 | .002 |
| R inferior parietal | 15 | 9 | HbO | 2.85 | .044 |
| R inferior parietal | 15 | 9 | HbR | -4.50 | .001 |
| R inferior parietal | 15 | 11 | HbO | 4.31 | .001 |

Note: Each channel corresponds to a pair of a source and a detector. Source: the light source number; Detector: the light detector number; HbO: oxyhaemoglobin; HbR: deoxyhemoglobin

### 6.3 Mathematics vocabulary in L2

The brain activation significantly increased in the left medial orbital frontal and middle frontal gyri, and the right inferior parietal gyrus during mathematics vocabulary in L2 (Table 4 and Fig. 3).

**Table 4.** Significant channels in the two contrasts.

| Region | Source | Detector | HbO/HbR | t | pFDR |
| --- | --- | --- | --- | --- | --- |
| **Mathematical vocabulary in L1 vs L2** |  |  |  |  |  |
| R superior frontal | 3 | 3 | HbO | 4.67 | <.001 |
| R middle frontal | 6 | 5 | HbO | 4.03 | .003 |
| **Mathematical vocabulary in L1 vs non-mathematical control in L1** |  |  |  |  |  |
| R superior frontal | 3 | 3 | HbO | 4.41 | .001 |

Note: Each channel corresponds to a pair of a source and a detector. Source: the light source number; Detector: the light detector number; HbO: oxyhaemoglobin.

### 6.4 Object recognition control in L1

The brain activation significantly increased in the left medial orbital frontal gyrus, and the right postcentral, inferior and superior parietal gyri, and precuneus during object recognition control in L1 (Table 3 and Fig. 3).

### 6.5 Mathematics Vocabulary in L1 vs L2

The results of the paired *t*-test revealed a significant difference in accuracy of children performing mathematics vocabulary in L1 and L2, *t(*29) = 1.71, *p* = 0.049, Cohen’s *d* = 0.31. Children had better performance in mathematics vocabulary in L1 (*M* = 0.86) than in L2 (*M* = 0.82). However, their RT during mathematics vocabulary in L1 and L2 showed no statistically significant difference, *t(*29) = 1.04, *p* = 0.153, Cohen’s *d* = 0.19.

The contrast of mathematics vocabulary in L1 vs L2 showed higher brain activation in the right superior and middle frontal gyri (Table 4 and Fig. 3). This contrast reveals underlying brain activation of processing mathematical vocabulary in different languages.

### 6.6 Mathematics Vocabulary vs Object Recognition in L1

The results of the paired *t*-test revealed no significant difference in accuracy of children performing mathematics vocabulary and object recognition in L1 *t(*30) = -0.15, *p* = 0.558, Cohen’s *d* = -0.03. However, their response times during mathematics vocabulary and object recognition in L1 showed a statistically significant difference, *t(*30) = 1.95, *p* = 0.030, Cohen’s *d* = 0.35. Children were slower during mathematics vocabulary in L1 (*M* = 1848) than object recognition in L1 (*M* = 1736).

The contrast of the mathematical vocabulary in L1 vs object recognition in L1 showed higher brain activation in the right superior frontal gyrus (Table 4 and Fig. 3). This contrast reveal underlying brain activation of mathematical processing.

### 6.7 Cognitive Measures

The results of the paired *t*-test comparing children’s performance in the mathematics language (MVT) in L1 (*M* = 22.07) and L2 (*M* = 21.17) showed no statistically significant difference, *t(*41) = 1.63, *p* = 0.112, Cohen’s *d* = 0.25. However, their performance in verbal short-term memory (digit span) in L2 (*M* = 3.55) was statistically better than in L1 (*M* = 3.00), *t(*41) = -3.20, *p* = 0.003, Cohen’s *d* = -0.49.

### 6.8 Correlations among Measurements

Correlations between all study measurements are presented in Table 5. Age did not show any statistically significant correlations with other variables. The number of siblings, however, was negatively correlated with math anxiety. Nonverbal IQ was positively correlated with listening comprehension, mathematics, and digit span in L2. Listening comprehension was strongly correlated with mathematics and positively correlated with digit span in L2, mathematics language in L1 as well as mathematics language in L2. Furthermore, mathematics exhibited significant positive correlation with digit span in L2, and mathematics language in L1 and L2. There was also a positive significant correlation between mathematics language in L1 and L2.

**Table 5.**
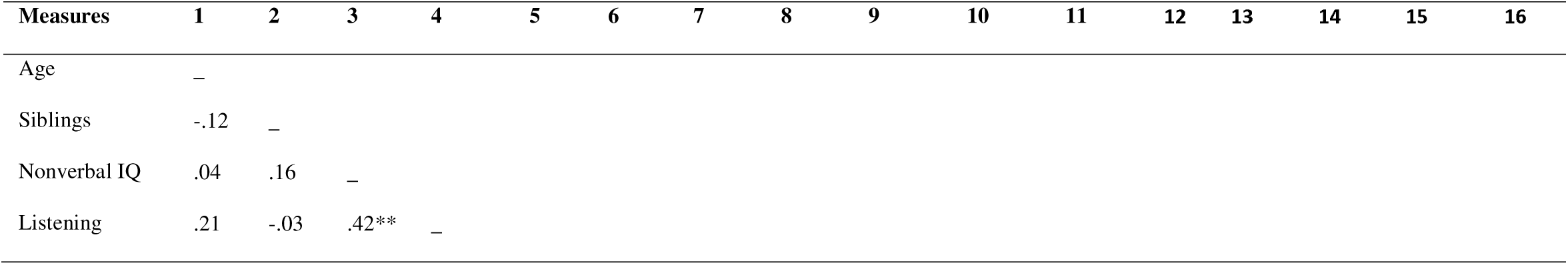
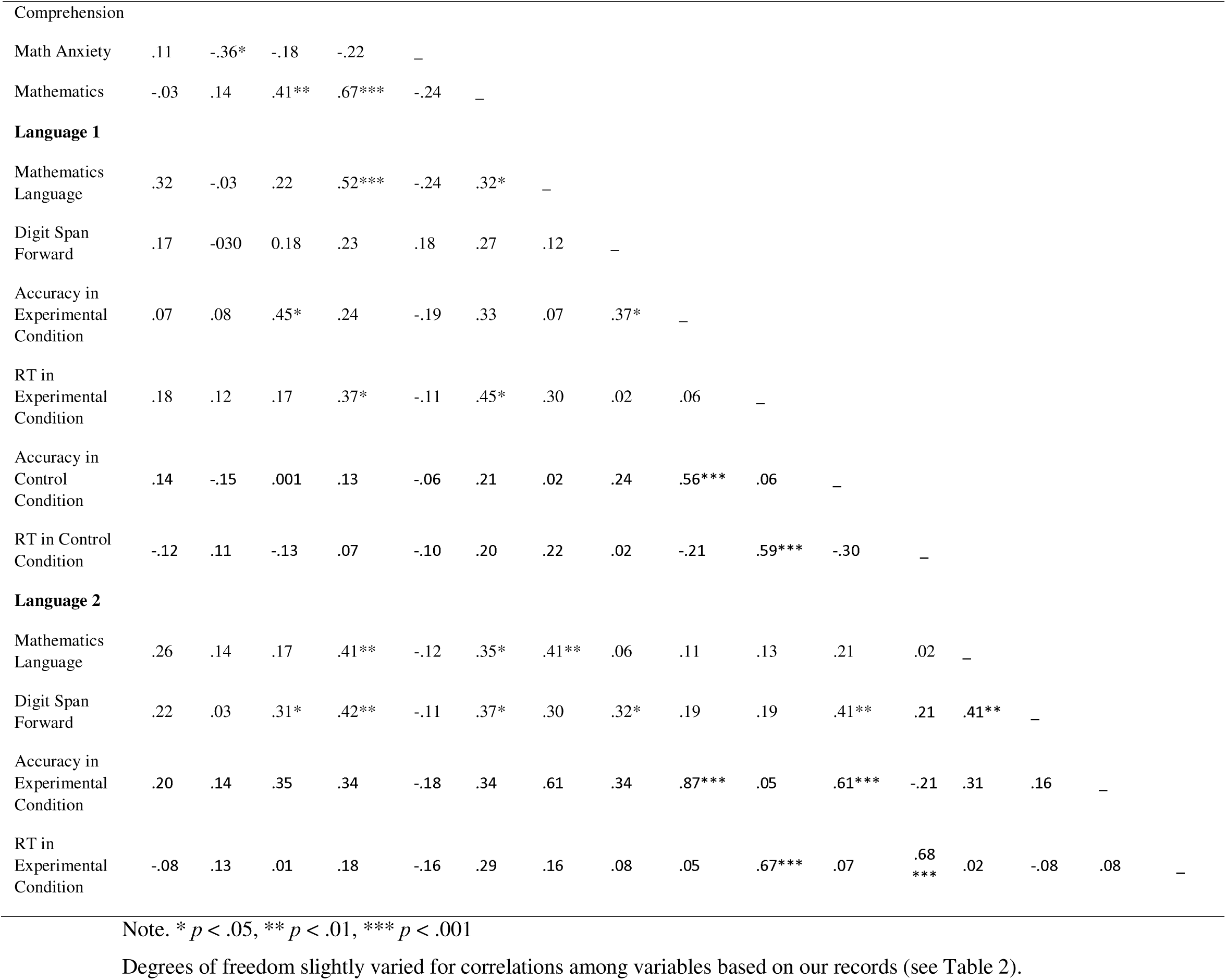
Correlations among Measurements.

| Measures | 1 | 2 | 3 | 4 | 5 | 6 | 7 | 8 | 9 | 10 | 11 | 12 | 13 | 14 | 15 | 16 |
| --- | --- | --- | --- | --- | --- | --- | --- | --- | --- | --- | --- | --- | --- | --- | --- | --- |
| Age | — |  |  |  |  |  |  |  |  |  |  |  |  |  |  |  |
| Siblings | -.12 | — |  |  |  |  |  |  |  |  |  |  |  |  |  |  |
| Nonverbal IQ | .04 | .16 | — |  |  |  |  |  |  |  |  |  |  |  |  |  |
| Listening | .21 | -.03 | .42** | — |  |  |  |  |  |  |  |  |  |  |  |  |
| Comprehension |  |  |  |  |  |  |  |  |  |  |  |  |  |  |  |  |
| Math Anxiety | .11 | -.36* | -.18 | -.22 | — |  |  |  |  |  |  |  |  |  |  |  |
| Mathematics | -.03 | .14 | .41** | .67*** | -.24 | — |  |  |  |  |  |  |  |  |  |  |
| Language 1 |  |  |  |  |  |  |  |  |  |  |  |  |  |  |  |  |
| Mathematics Language | .32 | -.03 | .22 | .52*** | -.24 | .32* | — |  |  |  |  |  |  |  |  |  |
| Digit Span Forward | .17 | -.030 | 0.18 | .23 | .18 | .27 | .12 | — |  |  |  |  |  |  |  |  |
| Accuracy in Experimental Condition | .07 | .08 | .45* | .24 | -.19 | .33 | .07 | .37* | — |  |  |  |  |  |  |  |
| RT in Experimental Condition | .18 | .12 | .17 | .37* | -.11 | .45* | .30 | .02 | .06 | — |  |  |  |  |  |  |
| Accuracy in Control Condition | .14 | -.15 | .001 | .13 | -.06 | .21 | .02 | .24 | .56*** | .06 | — |  |  |  |  |  |
| RT in Control Condition | -.12 | .11 | -.13 | .07 | -.10 | .20 | .22 | .02 | -.21 | .59*** | -.30 | — |  |  |  |  |
| Language 2 |  |  |  |  |  |  |  |  |  |  |  |  |  |  |  |  |
| Mathematics Language | .26 | .14 | .17 | .41** | -.12 | .35* | .41** | .06 | .11 | .13 | .21 | .02 | — |  |  |  |
| Digit Span Forward | .22 | .03 | .31* | .42** | -.11 | .37* | .30 | .32* | .19 | .19 | .41** | .21 | .41** | — |  |  |
| Accuracy in Experimental Condition | .20 | .14 | .35 | .34 | -.18 | .34 | .61 | .34 | .87*** | .05 | .61*** | -.21 | .31 | .16 | — |  |
| RT in Experimental Condition | -.08 | .13 | .01 | .18 | -.16 | .29 | .16 | .08 | .05 | .67*** | .07 | .68*** | .02 | -.08 | .08 | — |

Note. \* $p < .05$ , \*\* $p < .01$ , \*\*\* $p < .001$
Degrees of freedom slightly varied for correlations among variables based on our records (see Table 2).

Furthermore, accuracy of mathematics vocabulary performance in L1 showed a significant positive correlation with nonverbal IQ, digit span in L1 and accuracy of control condition and mathematics vocabulary performance in L2. Also, response times of mathematics vocabulary performance in L1 showed a positive significant association with listening comprehension, mathematics, and response times of mathematics vocabulary performance in L2. There was a significant positive strong correlation between response times of control condition L1 and both response times of mathematics vocabulary performance in L1 and L2.

## 7 Discussion

The aim of this study was to investigate the neural correlates of mathematics vocabulary processing (i.e., more and less) in L1 (i.e. Sesotho/isiZulu) and L2 (i.e. English) by using fNIRS in first graders in South Africa. We hypothesized (i) slower and less accurate responses, as well as higher activation in the prefrontal cortex when mathematics vocabulary is processed in L2 than in L1; and ii) slower and less accurate responses in mathematics vocabulary processing than object recognition, together with higher brain activation in the frontoparietal network in L1. In the first hypothesis, we did not expect a difference in parietal activation between L1 and L2 for mathematics vocabulary processing.

### 7.1 Mathematics vocabulary in L1 and L2

The results of our study show that higher accuracy in the L1 mathematics vocabulary task comes with the costs of higher cognitive demands in the right superior and middle frontal gyri for first graders who learn mathematics in L2. Although children responded, as hypothesized, more accurately in the L1 fNIRS mathematics vocabulary task, their response time did not differ. Contradictory to our hypothesis, children showed higher activation in the brain areas that support domain executive functions^27^ for the L1 mathematics vocabulary task. This result is in line with Sugiura et al.^28^ (2011), although in contradiction with others such as Linck et al.^58^ (2013). A possible explanation for higher activation in L1 mathematics vocabulary task is that children have richer phonological and semantic associations with vocabulary in L1^28^ and can integrate L1 vocabulary with mathematics concepts stored in the long-term memory^59^, leading to more cognitive control and integration in L1. Moreover, the young children in this study are at the learning stage of those concepts and are not yet masters. According to several neuroimaging studies of mathematics learning, they still heavily rely on domain-general cognitive resources for learning mathematical concepts. Therefore, higher frontal activation, that comes with better performance, during L1 mathematics vocabulary, provides further insights about mathematics learning and development in children.

The relationships between mathematics (PENS) and mathematics language, together with the differences between L1 and L2 mathematics vocabulary performance, lead us to conclude that although children may understand mathematical concepts, they may not yet be able to integrate L2 vocabulary with mathematics conceptual networks. In this study, isiZulu- or Sesotho-speaking learners have not yet automotized or integrated English mathematics vocabulary into a coherent mathematical conceptual network as much as L1 vocabulary. This raises the question whether children begin to use L1 mathematics vocabulary as mediator for the interpretation of L2 mathematics vocabulary as suggested by Wang et al.^25^ (2007). Further research is needed to investigate whether there exists a developmental shift in how L1 mediates L2 meaning in childhood compared to adulthood.

Further, children with better listening comprehension (in L1) also achieved higher scores in mathematics (PENS) and mathematics language (MVT) in L1 and L2. This means that children with better language comprehension may have a better understanding of mathematics concepts. In a multilingual setting like South Africa, it may mean that some children’s limited exposure to L2 may negatively affect their mathematics performance when measured in a different language than L1. The interpretations of these results are, however, influenced by many environmental factors^14^ such as young children’s varying exposure to mathematics vocabulary during their early years^19,22^ which makes it difficult to generalize the results to a larger linguistically diverse community.

### 7.2 Object recognition and mathematics vocabulary in L1

According to our hypothesis, mathematics vocabulary required longer RT than object recognition and a higher cognitive demand for supportive functions in the right superior frontal gyrus. However, contradictory to our hypothesis, there was no difference in their accuracy during these two tasks. As we expected, the mathematics vocabulary task has been more difficult for children than the object recognition tasks. Therefore, the more difficult task was completed slower and led to higher frontal activation, suggesting its demand for domain-general cognitive processes.

However, contradictory to our expectations, there was no parietal difference observed in the contrast of mathematics vocabulary and object recognition tasks (Fig. 3). While we observed right IPS activation during mathematics vocabulary in L1, object recognition in L1 led to a high activation in the right superior parietal lobule. The IPS, particularly the on the right hemisphere, is involved in processing magnitudes during almost any kind of numerical cognition task^61^, which is why mathematics vocabulary processing activates the right IPS. The right superior parietal lobe is related to visuospatial processing and supports spatial working memory (e.g., remembering object positions) which is why this area was activated in object recognition. These two activated regions subtracted each other in the contrast analysis resulting in no difference in the parietal region. One possible reason that the activation was subtracted, might be the low spatial resolution of fNIRS^37^. Note that more specific brain regions, like inferior temporal lobe, are involved in object recognition, however, due to our limitation, we were not able to record that region in the current study.

### 7.3 Exploratory findings

In addition to our priori hypotheses, we had two exploratory findings. Firstly, we observed a negative correlation between the number of siblings and mathematics anxiety. Although this suggests no causation, a possible interpretation is that children who have siblings that model confidence in mathematics or see siblings make mistakes and recover could experience lower mathematics anxiety. Interactions in home mathematics environments^62^ also create opportunities for siblings to engage in natural mathematics activities (e.g., splitting snacks, keeping score in games) that could make mathematics feel more practical and less intimidating. However, other environmental factors^14^ should be considered in the interpretation of these results and therefore further investigation is needed in future research.

Secondly, children had better performance in digit span in L2 than in L1, which was associated with other cognitive skills (i.e., listening comprehension, numeracy and similarity recognition). These findings can be explained by the practice of translanguaging^33,34^ in South Africa where most African speakers use English number words in their everyday discussions rather than number words in their home languages (e.g., isiZulu or Sesotho), by seamlessly switching between two known language structures. This is an example of how environmental factors influence concept development in specific populations and an example of the importance of direct research in majority countries^29^. However, this observation needs to be systematically explored in future research.

### 7.4 Challenges and limitations

Because this was one of the first fNIRS studies with children in South Africa, the main challenge of this study was to train isiZulu and Sesotho assistants to administer the fNIRS tasks. Eventually, we trained research assistants which led to a long period for data collection and because of inexperienced administrators, some of the data was lost. Another challenge in this study was the tradition of hair braiding that affects the contact between the fNIRS optodes and the childrens’ scalp, especially in the parietal regions. We could partially overcome this challenge by asking children not to braid their hair.

## 8 Conclusion

As one of the first educational neuroscientific studies in sub-Saharan countries, the fNIRS findings provided complementary insights about multilingual children’s mathematics learning at school entry. More specifically, it provided valuable insights in the neurocognitive mechanisms underlying mathematics vocabulary interpretation in different languages. The higher frontal activation during mathematics vocabulary suggests that young children are not yet automotized in the interpretation of mathematics vocabulary in L2. The multilingual context of South Africa, where children’s language of mathematics instruction is different to their home language may contribute to this result. This finding is not a generalization of the common studies of cognitive development in L1 and L2, where children acquire knowledge in their L1, but are exposed to L2 as well. In our sample, this is much more complex, because while South African children learn vocabularies in their home language (e.g., isiZulu), their “primary language or L1 for mathematics” is English. This is indeed the case of several other colonised sub-Saharan countries that their number system–or broadly speaking the academic language–has been contributed to be the coloniser’s language. In light of these results, together with Bleses et al.^1^ (2023) and Bezuidenhout et al.^42^ (2025), we suggest that teachers explicitly teach mathematics vocabulary, especially by using the known mathematics vocabulary in L1 during the preschool years as foundation to teach L2, preparing children for the interpretation of mathematics vocabulary in L2 in formal mathematics instruction.

## Acknowledgements

We would like to thank all participating children, parents, and the school for participating in this study. We would also thank our research assistants, Rhulani Clayton Ramasodi, Ellen Tebello Mololo, Nozipho Motolo, Thabitha Mapuleng Tsepesi and Zongezile Bright Zenzile for supporting data collection.

## Authors’ contribution

Bezuidenhout, Soltanlou and Henning conceptualised and designed the study; Bezuidenhout and assistants collected data; Nemati, Borjkhani and Soltanlou analysed and interpreted the data; all authors drafted and reviewed the manuscript.

## Data Availability

The data that support the findings of this study are available on request from the corresponding author.

